# Characterisation of PHA synthase (PhaC) from the thermophilic bacterium *Caldimonas thermodepolymerans* DSM15344

**DOI:** 10.64898/2026.09.18.752759

**Authors:** H. Majerová, M. Benešík, J. Krouská, P. Sedláček, P. Dvořák

**Affiliations:** Department of Experimental Biology, Faculty of Science, Masaryk University, Brno, Czech Republic; Institute of Physical and Applied Chemistry, Faculty of Chemistry, Brno University of Technology, Brno, Czech Republic

## Abstract

Polyhydroxyalkanoates (PHAs) are microbial polyesters that represent a sustainable alternative to fossil fuel-based plastics. These biodegradable polymers have properties similar to conventional plastics and can be produced from waste materials. The key enzyme in PHA synthesis is PHA synthase (PhaC) and despite its importance, only a few PhaCs from extremophilic microorganisms have been characterised in detail. Here we characterised PHA synthase enzyme from the thermophilic bacterium *Caldimonas thermodepolymerans* DSM15344, an attractive candidate for PHA production from lignocellulosic residues due to its unique properties such as utilisation of xylose-rich substrates and high PHA yields on this pentose sugar. PhaC was successfully expressed in the surrogate host *E. coli* BL21 (DE3) GOLD, purified to homogeneity and its enzymatic activity was confirmed *in vitro*. The optimal pH range and temperature were determined as 6.3–6.9 and 50 °C, respectively. The melting temperature of PhaC was detected at 57.83 ± 0.04 °C and the enzyme storage stability was assessed. The tertiary structure, including the enzyme active site, was predicted using available bioinformatic tools and the predicted catalytic residues were supported by site-directed mutagenesis. This study represents the first biochemical characterisation of PHA synthase from *C. thermodepolymerans*, expanding the limited set of characterised thermophilic PhaC enzymes.

## Introduction

In our current world, plastic materials represent a necessity and opportunity, but also a global problem due to pollution and their production from fossil fuels. Only about 9% of plastic waste is recycled and even if we recycled it more efficiently, the problems of microplastic pollution and plastic production from non-renewable resources would still be present (Lim 2021; Houssini et al. 2025). Therefore, a sustainable alternative is needed. One of the promising alternatives are polyhydroxyalkanoates, polymers of microbial origin that are renewable, biodegradable and biocompatible. They are currently used in various applications, including cosmetics, packaging or medicine (Meereboer et al. 2020; Park et al. 2024). The material properties of PHA depend on monomers incorporated into the polyester, which allow production of materials with tailored properties (Koller 2019). Despite these advantages, they represent only a negligible fraction of produced plastics. The main reasons for low industrial production of PHA are high cost and non-ideal properties of the most abundant PHA – poly(3-hydroxybutyrate) (Zytner et al. 2023). However, these problems can be solved by engineering of PHA producers or by implementation of Next Generation Industrial Biotechnology (NGIB). This concept uses extremophilic microorganisms for biotechnological production and can reduce the price of the final polymer (Chen and Jiang 2018). Today, mainly halophiles are being used in the NGIB concept, such as engineered *Halomonas bluephagenensis* (Fu et al. 2014; Obruča et al. 2022). In contrast to PHA production in halophiles, thermophilic PHA producers are not deeply investigated, although they bring many advantages. Biotechnological production in thermophiles is less susceptible to mesophilic contamination and could save energy on sterilisation or cooling of the bioreactor (Ibrahim and Steinbüchel 2010; Bhalla et al. 2013; Obruča et al. 2022).

Thermophilic bacterium *Caldimonas thermodepolymerans* is an attractive candidate for the NGIB concept. This bacterium can produce high yields of PHA (up to 87% CDW) from xylose, at its optimal temperature 50 °C (Kouřilová et al. 2020; Zhou et al. 2023). Besides xylose, this bacterium can grow on other lignocellulosic sugars such as cellobiose, has been recently engineered to grow on glucose (Hajkova et al. 2026), and is fairly resistant to inhibitors present in lignocellulosic hydrolysates, such as furfural, gallic or levulinic acid (Kouřilová et al. 2020; Kouřilová et al. 2021). As substrates represent a significant part of the cost, use of waste lignocellulose brings not only environmental but also economic advantage (Kourmentza et al. 2017). Besides reducing production cost, it is important to ensure the production of PHA with desired material properties.

The type of PHA produced depends on the metabolic capabilities of the microorganism and the characteristics of its PHA synthase (PhaC). PhaC is the key enzyme of PHA biosynthesis and catalyses the final step – polymerisation of various (R)-hydroxyacyl-CoA thioesters into PHA (Koller 2019). Understanding these enzymes is therefore essential for improving PHA production and enabling protein engineering approaches to advance large-scale production and applications of PHA. Currently, four classes of PhaCs are distinguished, which differ in the length of preferred substrates and the type of the enzyme subunits. Classes I, III, and IV PhaC usually produce short-chain-length PHA, while class II typically produces medium-chain-length PHA (Rehm 2003). Among them, class I PhaC is the most researched one, with several crystal structures determined (Zher Neoh et al. 2022; Chek et al. 2025). Characterised PhaCs are mainly from mesophilic or halophilic microorganisms, such as *Cupriavidus necator* or *Haloferax mediterranei* (Han et al. 2010; Kim et al. 2017a). In contrast, only few thermophilic PhaC has been described, e.g. PHA synthases from *Thermus thermophilus* or *Cupriavidus* sp. S-6 (Pantazaki et al. 2003; Sheu et al. 2012). However, in a recent publication, no evidence of polyhydroxyalkanoate production by *T. thermophilus* was found (Schroyen et al. 2025).

In this study, we characterise class I PhaC from thermophilic bacterium *C. thermodepolymerans*. The *phaC* gene was cloned and expressed in a surrogate host *E. coli* BL21 (DE3) GOLD and, after purification, characterised using biochemical and biophysical methods. Its structure was assessed by various bioinformatic tools and its predicted catalytic triad was confirmed by mutagenesis experiments. This work represents one of the first experimentally characterised PhaC enzymes from a thermophilic bacterium.

## Methods

### Bacterial strains and culture conditions

The bacterial strains *Escherichia coli* CC118 (Manoil and Beckwith 1985) and *E. coli* BL21(DE3) GOLD (Agilent Technologies) were routinely grown in 50 ml of lysogeny broth (LB; Serva) in shake flasks or on agar plates (1.6% w/v) at 37 °C. Ampicillin was added to the medium at a final concentration of 150 µg.ml^-1^.

### Bioinformatic analysis

The theoretical molecular weight of the protein was calculated using ProtParam (Expasy). DNA and protein sequences were analysed in Benchling. To identify catalytic residues, a multiple sequence alignment (MSA) of PhaC homologues with experimentally determined structures was performed using Clustal Omega (Madeira et al. 2024). The alignment included protein sequences from *C. thermodepolymerans* DSM15344 (GenBank: PPE70723.1), *Cupriavidus necator* H16 (CAJ92572.1) and *Chromobacterium* sp. USM2 (ADL70203.1) and visualised using the ESPript 3.2 (Gouet et al. 1999). The secondary structure scheme was generated by PSIPRED server (McGuffin et al. 2000) and the tertiary structure was predicted using AlphaFold Server (AlphaFold 3) (Abramson et al. 2024).

### Gene synthesis and general cloning procedures

The *phaC* gene (locus tag IS481_08630) from *C. thermodepolymerans* DSM15344 was codon optimised for expression in *E. coli* and synthesized by the company Azenta. The synthesized gene was cloned into pET-21b, with a 6x His tag sequence at the gene’s 3’ terminus, by restriction cloning. The annotated pET-21b plasmid containing the *phaC* gene is shown in Fig. S1. All plasmids used in this study are listed in Table 1. Plasmid DNA was isolated using the E.Z.N.A. Plasmid DNA Mini Kit I. DNA purity and concentration were determined by NanoDrop 2000 (Thermo Fisher Scientific).

**Table 1.**
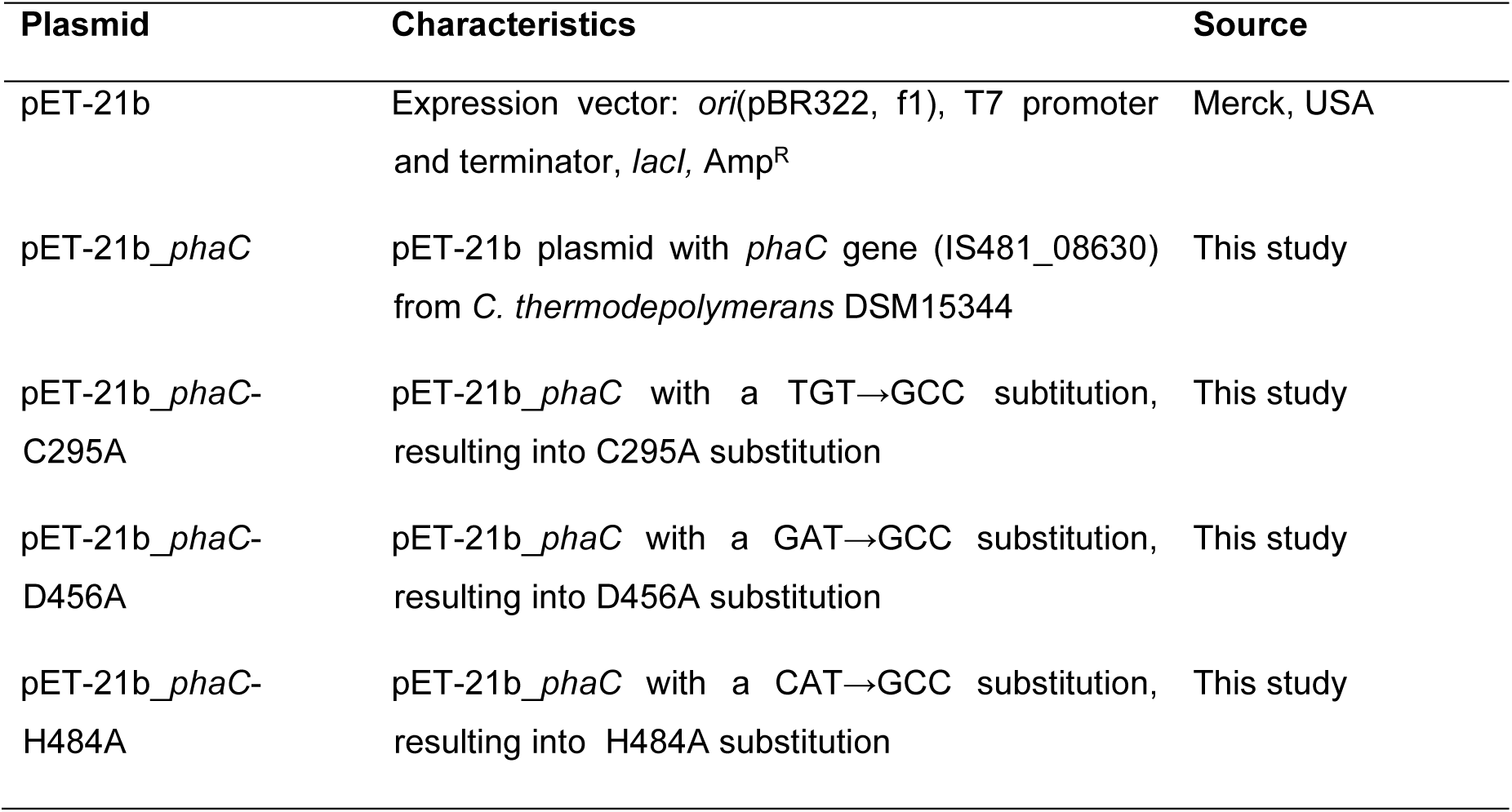
Plasmids used in this study.

| Plasmid | Characteristics | Source |
| --- | --- | --- |
| pET-21b | Expression vector: *ori*(pBR322, f1), T7 promoter and terminator, *lacI*, Amp<sup>R</sup> | Merck, USA |
| pET-21b_phaC | pET-21b plasmid with *phaC* gene (IS481_08630) from *C. thermodepolymerans* DSM15344 | This study |
| pET-21b_phaC-C295A | pET-21b_phaC with a TGT→GCC substitution, resulting into C295A substitution | This study |
| pET-21b_phaC-D456A | pET-21b_phaC with a GAT→GCC substitution, resulting into D456A substitution | This study |
| pET-21b_phaC-H484A | pET-21b_phaC with a CAT→GCC substitution, resulting into H484A substitution | This study |

### Construction of *phaC* mutant variants

PhaC variants with mutated catalytic triad residues (pET-21b_*phaC*-C295A, pET-21b_*phaC*-D456A and pET-21b_*phaC*-H484A) were prepared by replacing the codon encoding the respective catalytic amino acid with an alanine-encoding codon (GCC). The site-directed mutagenesis was performed by USER cloning using mutagenic primers. Primers (Table S1) containing deoxyuridine residues were designed using the AMUSER online tool (Bitinaite et al. 2007; Genee et al. 2015) and their annealing temperatures were calculated by the NEB T*_m_* Calculator. DNA was amplified by Q5U Hot Start High-Fidelity DNA Polymerase (New England Biolabs; NEB) according to the manufacturer’s protocol. Amplified DNA was digested with the DpnI enzyme (NEB) for 30 min at 37 °C and subsequently treated with USER enzyme (NEB). USER reaction mixture (100 ng of each vector and insert, 1 µl of T4 ligase buffer, 1 µl of USER enzyme, Milli-Q H_2_O up to 12 µl) was incubated in a heat block at 37 °C for 25 min and then at 25 °C for a further 25 min. To achieve covalent binding, 1 µl of T4 ligase (NEB) was added to the samples and incubated for 40 minutes at room temperature. Constructed plasmids were transformed into chemically competent *E. coli* CC118 by the heat shock method. Transformants were selected on LB agar plates with ampicillin and the presence of the desired mutation was verified by Sanger sequencing (SEQme, Czech Republic). Confirmed transformants were stored in LB medium with 20% (v/v) sterile glycerol at −70 °C.

### PhaC expression and purification

The appropriate plasmid was transformed into the expression strain *E. coli* BL21(DE3) GOLD and was grown at 37 °C and 200 rpm (N-Biotek shaker) to an OD_600_ ≈ 0.5. Expression was induced with IPTG at a final concentration of 0.4 mM. After 4 hours of cultivation at 30 °C, the cells were harvested by centrifugation at 4,500 rpm (Universal 320R; Hettich) for 15 minutes at 6 °C and the biomass was stored at −20 °C.

For purification, cells were resuspended in sonication buffer (50 mM Tris, 20 mM imidazole, 200 mM NaCl, 0.1% Triton X100, pH 7.5) and disrupted by sonication at 80% amplitude (NexTgen Lab120; SinapTec) with repeated cycles of 1 min pulses and 1 min pauses, while kept on ice. Cell lysate was centrifuged at 24,000 g (Avanti J-26S XPI; Beckman Coulter) for 20 minutes at 10 °C and the supernatant was purified by nickel affinity chromatography using a FPLC system (NGC Chromatography System; Bio-Rad). The cell-free extract was loaded onto a HisTrap HP column (Cytiva) equilibrated with Tris buffer (50 mM Tris, 20 mM imidazole, 200 mM NaCl, pH 7.5). Bound proteins were eluted using an increasing imidazole gradient (20–500 mM). The target protein was transferred into an appropriate buffer using an Amicon Ultra-4 Centrifugal Filter Unit (30 kDa; Millipore). Independent protein preparations were used for separate characterisation experiments.

Protein concentration was determined using the Bradford assay with Bradford reagent (Sigma-Aldrich), with bovine serum albumin (BSA) as a standard. SDS-PAGE was performed using a 12% resolving gel and 4% stacking gel. Protein samples were mixed with 4x sample loading buffer (final concentration 1x) and boiled at 95 °C for 5 minutes. Wells of the gel were loaded with 12.5 µL of boiled sample and 5 µL PageRuler Prestained Protein Ladder (10 to 180 kDa) was used as a marker. Electrophoresis was performed at 80 V for approximately 80 minutes. The gel was then washed in distilled water for 10 minutes and stained with Quick Coomassie stain for 1 hour. The gel was then rinsed several times, destained overnight in distilled water and photographed the following day.

### PhaC activity assay

PhaC activity was determined by measuring free coenzyme A (CoA) released during PHA polymerisation. Free CoA was detected by reaction with 5,5′-dithiobis(2-nitrobenzoic acid) (DTNB) (Ellman 1959; De Roo et al. 2000). The reaction mixture (250 µl total) contained 200 µl DTNB (1 mM), 25 µl 3-hydroxybutyryl-CoA (2 mM) and 25 µl enzyme (0.5 mg.ml^−1^ PhaC). Measurements were carried out in a 96-well plate (BRAND 96-well pureGrade, polystyrene plates) at 50 °C using a BioTek Microplate Reader (Agilent). For the negative control, PhaC was replaced with 0.5 mg.ml^−1^ BSA. Absorbance was measured at 412 nm at 1 min intervals and enzyme activity was calculated from the slope of the linear phase of the absorbance-time curve (ΔA/min). The amount of CoA released was quantified using a calibration curve prepared from appropriate dilutions of 0.2 mM free CoA in 1 mM DTNB. PhaC activity was represented either as specific or relative activity. All reagents were prepared in Tris buffer (50 mM Tris, 150 mM NaCl, pH 6.9 at 50 °C) unless specified otherwise.

### Determination of pH and temperature optima

To determine the pH optimum of PhaC, an activity assay was performed with the following modifications. DTNB was diluted in either a citrate buffer (50 mM citrate, 150 mM NaCl; in a pH range 5.4 – 6.7) or a Tris buffer (50 mM Tris, 150 mM NaCl; in a pH range 6.9 – 7.9), adjusted to the appropriate pH. The temperature optimum was determined by performing activity measurements at different temperatures. To eliminate the influence of the pH-temperature dependency, the pH of the Tris buffers was adjusted at the corresponding temperatures. For each temperature, 1 mM DTNB was prepared using the appropriate Tris buffer.

### Differential scanning calorimetry

For differential scanning calorimetry (DSC), purified PhaC (1 mg.ml^−1^) in Tris buffer (50 mM Tris, 150 mM NaCl) was used. Thermal scans were obtained using a MicroCal PEAQ-DSC system, within the temperature range of 40–90 °C. Tris buffer was used as a blank in the reference capillary cell during the measurement. Baseline subtraction was performed in MicroCal PEAQ-DSC software. DSC was performed in three independent replicates.

### Storage stability assessment

To evaluate PhaC storage stability, enzyme activity was measured at optimised conditions, after purification (time zero) and after 30 days of storage. The purified protein was divided into four aliquots, which were stored under the following conditions: at 23 °C, 4 °C, −20 °C and −60 °C. For samples stored at −20 °C and −60 °C, glycerol was added to a final concentration of 20%. To account for the effect of glycerol on enzyme activity, PhaC activity at time zero was measured both in the presence and absence of glycerol (final concentration in the reaction mixture was 0.4%). Relative activity was normalised to the corresponding initial activity at time zero (set as 100%), measured either in the presence or absence of glycerol.

### Data analysis

Data are presented as mean values with corresponding standard deviations (SD). The number of biological (n) or technical replicates is specified in the corresponding figure legends. When indicated, statistical significance was assessed using a two-tailed Student’s *t*-test in Microsoft Office Excel (Microsoft).

## Results

### The *phaC* gene cloning, heterologous expression and purification of the PhaC protein

Previously reported genome sequencing of *Caldimonas thermodepolymerans* DSM15344 revealed one gene encoding PHA synthase (PhaC; locus tag IS481_08630) (Musilová et al. 2023). The identified *phaC* gene is 1,698 bp long and the resulting protein consists of 565 amino acids. To allow easy manipulation and production, the *phaC* gene was synthesized, cloned into the expression plasmid pET-21b (Table 1), and expressed in the surrogate host *E. coli* BL21(DE3) GOLD. Successful production of PhaC and subsequent purification (Fig. S2) were visualised by SDS-PAGE, revealing a sole band corresponding to PhaC purified to homogeneity (62.7 kDa; Fig. 1).

**Fig. 1.**
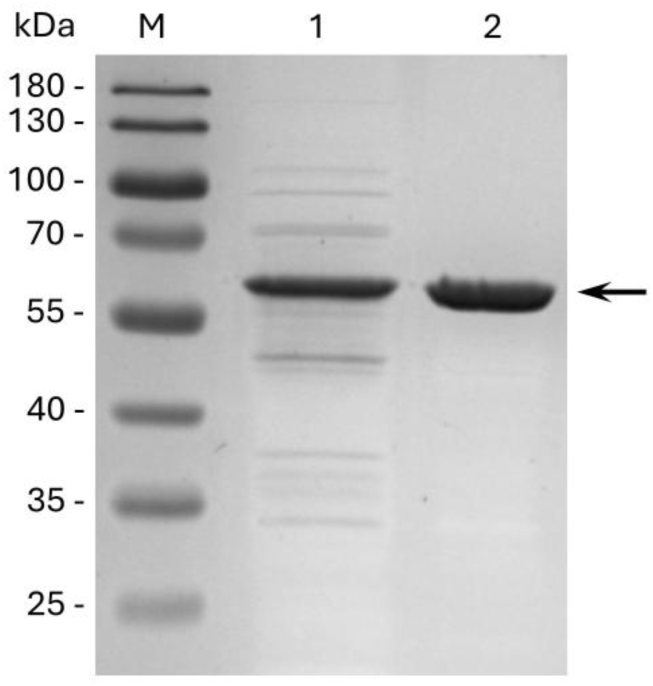
**SDS-PAGE of PhaC samples before and after purification**The samples were separated on a 12% SDS-PAGE gel and stained with Quick Coomassie Stain. M: PageRuler Prestained Protein Ladder (10 to 180 kDa); lane 1: cell-free extract; lane 2: purified PhaC (marked with an arrow) after Ni-NTA affinity chromatography.

To verify that the purified PhaC was enzymatically active, its activity was measured by an assay based on Ellman’s reaction (Ellman 1959; De Roo et al. 2000). The initial activity measurement was carried out at 50 °C and confirmed that purified recombinant PhaC retained its enzymatic activity (Fig. S3). An initial four-minute lag phase was observed and the specific activity was determined to be 0.327 ± 0.023 U.mg^−1^.

### Effect of pH and temperature on PhaC activity

For the determination of the pH optimum, PhaC activity was measured in citrate and Tris buffers at 50 °C and varying pH values (Fig. 2a). The highest activity was observed at pH 6.7 in citrate buffer and the enzyme also retained high activity at pH 6.3 (citrate buffer) and at pH 6.9 (Tris buffer). In contrast, at pH 5.8 and pH 7.4, PhaC lost more than 50% of its activity. Following the determination of the pH optimum, the temperature optimum was assessed in Tris buffer at pH 6.9. Even though the highest activity was not achieved at this pH, we decided to continue all measurements in Tris buffer, to maintain consistency with previous assays. To account for the temperature dependence of Tris buffer, pH was adjusted at each corresponding temperature (Durst and Staples 1972). Relative activity increased with temperature, reaching the maximum at 50 °C (Fig. 2b). At higher temperatures of 55 °C and 60 °C, activity decreased by approximately 40% and 50%, respectively.

**Fig. 2.**
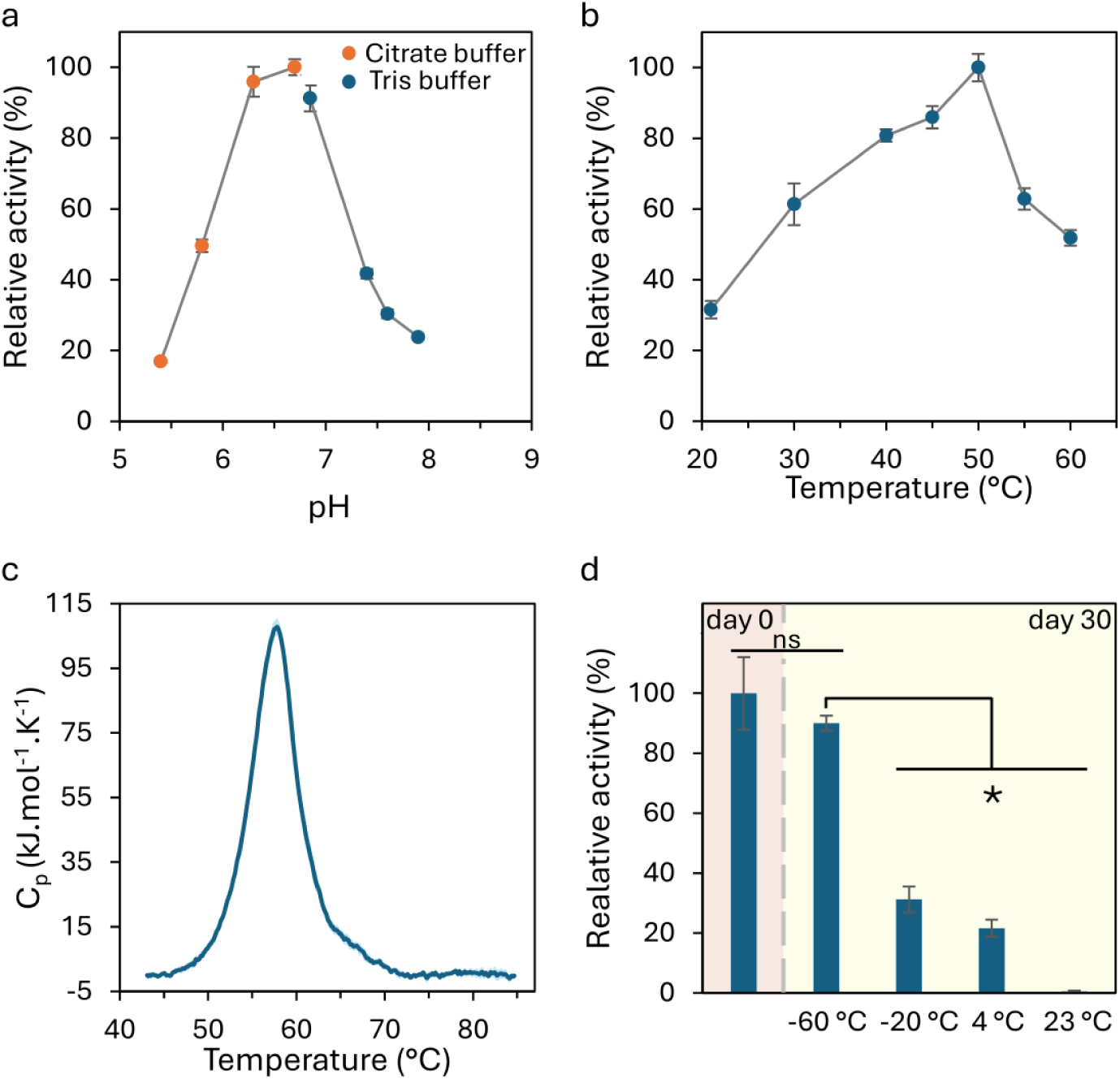
Determination of pH and temperature optima and storage stability of PhaC. **(a)** Dependence of PHA synthase relative activity on pH was measured at 50 °C. Orange dots represent measurements performed with citrate buffer (50 mM citrate, 150 mM NaCl), blue dots represent measurements with Tris buffer (50 mM Tris, 150 mM NaCl). **(b)** Temperature optimum of PhaC was measured in Tris buffer at pH 6.9. **(c)** Thermogram of PhaC (1 mg.ml^−1^ in 50 mM Tris, 150 mM NaCl) showing a single, sharp endothermic unfolding transition with slight asymmetry. The y-axis represents molar heat capacity (C_p_). **(d)** Storage stability of PhaC at different conditions. Day 0 – activity measured after purification set as 100% relative activity. Day 30 – relative activities of samples after a month of storage at the appropriate temperatures. The bars lined together and labelled ’ns’ are not significantly different; statistical significance (P < 0.01) is marked with an asterisk. For the determination of the P value, Student’s t-test was used. All data points represent the mean ± standard deviation (SD) of three biological replicates (n=3).

### Thermal and storage stability of PhaC

Thermal stability of PhaC was analysed by differential scanning calorimetry (DSC). The resulting thermogram showed a single endothermic transition corresponding to the enzyme unfolding, starting approximately at 46 °C (Fig. 2c), which is below the temperature optimum determined by activity measurements. The maximum of the heat capacity curve representing the melting temperature (*T*_m_) of the protein was observed at 57.83 ± 0.04 °C.

The storage stability of PhaC was evaluated after 30 days of storage in four different conditions (Fig. 2d). Activity measured after purification was used as a time zero reference. The presence of glycerol, which was added to samples stored in the freezer, significantly reduced PhaC activity. Specifically, the final concentration of 0.4% glycerol in the reaction mixture resulted in an approximately 50% decrease in activity compared to the glycerol-free sample. After 30 days of storage, enzyme activity was strongly influenced by storage conditions. The sample stored at −60 °C retained the highest activity, approximately 90% relative activity. In contrast, PhaC stored at −20 °C retained only about 30% of its initial activity. Similarly, storage at 4 °C resulted in approximately 20% remaining activity, while the sample stored at 23 °C was completely inactive.

### PhaC structure

The predicted secondary structure revealed that PhaC consists mainly of α-helices, except for the core of the C-terminal domain, which forms an α/β-hydrolase-like structure (Fig. S4). Structural features were identified using multiple sequence alignment (MSA; Fig. S5) by comparison with homologous enzymes whose structures have been previously described. PhaC is organised into N- and C-terminal domains (Fig. 3b). The C-terminal domain consists of an α/β core subdomain and a CAP subdomain with a LID region. The predicted catalytic residues C295, D456, and H484 were all located in the C-terminal domain.

**Fig. 3.**
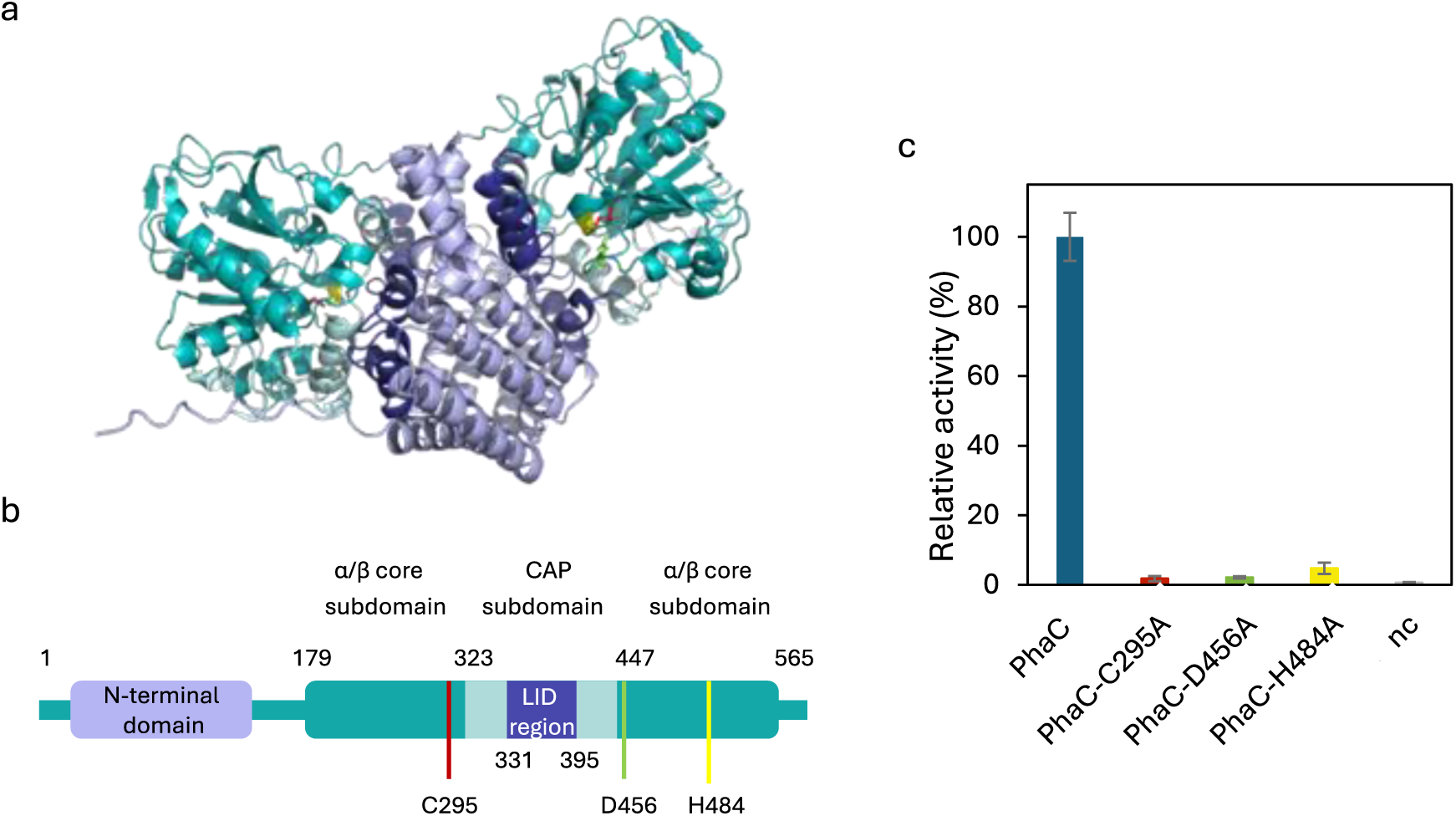
Prediction of PhaC catalytic residues and their verification by mutagenesis. **(a)** Dimeric structure of PhaC predicted using AlphaFold 3. The N-terminal domain is shown in light blue. The C-terminal domain consists of α/β core subdomain, CAP subdomain and LID region, shown in teal, light green and dark blue, respectively. The predicted catalytic residues C295, D456 and H484 are shown in red, green and yellow. **(b)** Schematic representation of monomeric PHA synthase with N- and C-terminal domains. The domain responsible for enzyme activity is the C-terminal domain, which contains α/β core subdomain and CAP subdomain with a LID region. The α/β core subdomain ranges from P179 to T323 and from P447 to A565. The CAP subdomain ranges from T323 to P447. The catalytic triad is located in α/β core subdomain. **(c)** Activity assay with PhaC mutated variants. WT PhaC activity was set as 100% relative activity; PhaC mutants PhaC-C295A, PhaC-D456A and PhaC-H484A reached 1.8%, 2.1% and 4.8% of the activity of WT enzyme, respectively.

The tertiary structure of PhaC from *C. thermodepolymerans* has not been crystallised so far, and its predicted structure is currently available in the Uniprot database (A0A2S5T6U4). As PhaC was confirmed to exist predominantly as a dimer (Wodzinska et al. 1996; Chek et al. 2025) and the model in the database is available only in monomeric form, its tertiary structure was predicted in the dimeric form (Fig. 3a). The AlphaFold evaluation scores were 0.15 for the interface predicted template modelling (ipTM) and 0.43 for the predicted template modelling (pTM; scores range from 0 to 1, with higher values indicating higher confidence in the protein model).

### Verification of PhaC catalytic triad residues

To verify the catalytic triad predicted by MSA, three plasmid constructs with mutated *phaC* gene were prepared by site-directed mutagenesis. In each construct, the codon encoding one of the catalytic residues was substituted with alanine encoding codon, PhaC-C295A, PhaC-D456A and PhaC-H484A. The presence of modified codons was verified by Sanger sequencing (Fig. S6), followed by expression and purification of PhaC variants (Fig. S7 and Fig. S8). The effect of the mutations was assessed by an activity assay. In all three mutants a loss of activity was observed in comparison to wild-type (WT) PhaC, supporting the proposed catalytic role of these amino acid residues (Fig. 3c).

## Discussion

Detailed insight into PHA synthases, the key enzymes of PHA synthesis, is necessary to understand PHA production. To date, there are PhaC characterised from many microorganisms, with one of the most researched PhaC from *C. necator* (Zhang et al. 2000; Kim et al. 2017a; Zher Neoh et al. 2022). However, there is a lack of data on these enzymes from the thermophilic microorganisms (Schroyen et al. 2025). Here we characterise PhaC from moderate thermophile *C. thermodepolymerans*. For experimental characterisation, the *phaC* gene from *C. thermodepolymerans* DSM15344 was synthesised, heterologously expressed in *E. coli* BL21(DE3) GOLD, and purified. The enzymatic activity of the recombinant PhaC was confirmed by an activity assay performed at 50 °C. This temperature was chosen as it is the optimal growth temperature of *C. thermodepolymerans* and therefore probably close to the optimal temperature of PhaC (Zhou et al. 2023; Grybchuk-Ieremenko et al. 2025). At the beginning of the activity measurement a lag phase was observed, a phenomenon common in PHA synthases and a lag phase of similar length was observed, e.g. in PhaC from *Chromobacterium* sp. USM2 (Jia et al. 2016). The length of the lag phase can vary between PhaCs, ranging from less than a minute up to 20 minutes and its origin remains unclear (Wodzinska et al. 1996; Zhang et al. 2000). One possible explanation is a requirement for PhaC dimerization, as longer lag phases were observed at lower protein concentrations (Gerngross et al. 1994; Kim et al. 2017b). Another possibility involves local structural changes in the tunnel due to interactions with the nascent PHA oligomer (Chek et al. 2025).

For further PhaC characterisation and optimisation of the activity assay, pH and temperature optima were determined. PhaC pH optimum was determined to be pH 6.3–6.9, with major activity decrease outside this range (Fig. 2a). Zhou et al. reported the highest yields of PHA at pH 7 and 8 in *C. thermodepolymerans*. This slightly more acidic pH optimum observed for PhaC may reflect the contribution of other enzymes involved in PHA biosynthesis (Zhou et al. 2023). A similarly narrow pH optimum was observed in PhaC from *C. necator*, between pH 6.8 and 7.2 (Zhang et al. 2000). The temperature optimum was determined at pH 6.9 in Tris buffer. PhaC activity increased with temperature, reaching a maximum at 50 °C (Fig. 2b), consistent with the optimal temperature for PHA production in *C. thermodepolymerans* (Zhou et al. 2023). At higher temperatures, a major decrease in activity was observed, probably caused by the thermal denaturation of the enzyme, as the melting temperature of PhaC determined by DSC was ∼58 °C (Fig. 2c). The heat capacity begins to increase around 46 °C, suggesting that PhaC may start to denature below its optimal temperature, but its activity remains high enough to outweigh this. The difference between T*_m_* and optimal temperature is approximately 8 °C, which is common for many mesophilic and thermophilic enzymes (Gault et al. 2021). The DSC revealed a single endothermic transition, indicating mostly cooperative unfolding of PhaC. The peak was sharp but slightly asymmetric, possibly due to the aggregation of the unfolded enzyme.

The influence of storage conditions on PhaC activity was evaluated by a 30 day storage stability experiment. Even at a low final concentration of 0.4%, glycerol significantly inhibited PhaC activity, suggesting that PhaC is unusually sensitive to glycerol. This may be due to glycerol influence on enzyme conformation or active–site accessibility. After 30 days, storage temperature strongly influenced enzyme stability. Storage at −60 °C effectively preserved PhaC activity, while storage at higher temperatures resulted in substantial activity loss and the sample stored at 23 °C was inactive (Fig. 2d). The large loss of activity at −20 °C may reflect insufficient protection by glycerol at a lower concentration (20%) during slow freezing. This can cause ice crystal formation and protein unfolding or aggregation. In contrast, rapid freezing at −60 °C, minimizes these effects (Cao et al. 2003).To optimise storage conditions of PhaC, glycerol concentration must be carefully balanced to maintain enzyme stability while minimizing inhibitory effects. A possible strategy is to store PhaC at higher protein concentrations, so glycerol would be more diluted in the reaction mixture. Further studies could investigate long-term stability and the impact of repeated rethawing on PhaC activity.

Structural analysis showed that PhaC consists predominantly of α-helices with a central α/β-hydrolase structure, which is characteristic for PHA synthases (Zher Neoh et al. 2022). The AlphaFold evaluation scores (ipTM: 0.15, pTM: 0.43) indicate low confidence in the global fold and inter-chain interactions, suggesting that while the PhaC monomers may adopt plausible conformations, their orientation and interaction are likely incorrect. This is supported by low local scores observed in the N-terminal domain and the LID region. As these regions are flexible and their conformational changes are necessary for enzyme activity and dimerization, their low confidence affects the dimer interface (Chek et al. 2020; Zher Neoh et al. 2022). Overall, the results suggest that the PhaC model is only partially reliable and further modelling is required to obtain a more accurate PhaC dimer structure.

The catalytic residues predicted by MSA correspond to the catalytic triad, which is conserved among class I PhaCs, reported for *C. necator* H16 and *Chromobacterium* sp. USM2 (Wittenborn et al. 2016; Chek et al. 2017). To verify the predicted catalytic triad, plasmid constructs carrying mutated *phaC* gene variants were prepared using USER cloning, expressed, purified and their activity was measured alongside the WT PhaC under identical conditions. The activity of all mutants was negligible compared to the WT (Fig. 3b). This indicates that residues C295, D456 and H484 are necessary for PhaC activity and likely form the catalytic triad. This result is also consistent with our prediction and previously studied PhaCs (Wittenborn et al. 2016; Chek et al. 2017). To rule out the possibility that the observed loss of activity resulted from structural alterations induced by the mutation, the secondary structures of WT PhaC and corresponding mutants will be assessed by circular dichroism spectroscopy.

In conclusion, this study provides the first experimental characterisation of PhaC from the thermophilic bacterium. *C. thermodepolymerans*. The enzyme was successfully produced and showed highest activity at 50 °C, consistent with the optimal growth temperature of its host, and pH 6.3-6.9. PhaC also showed notable sensitivity to storage conditions, especially to glycerol. Bioinformatic analysis was used to visualise enzyme structure and predict its catalytic triad, whose role was supported by a mutagenesis experiment. Overall, these results establish the key biochemical properties of PhaC from *C. thermodepolymerans* and provide a basis for further studies and potential engineering of this enzyme.

## Author Contributions

**Hana Majerová**: Conceptualization; investigation; methodology; writing – original draft; writing – review and editing; visualization; formal analysis. **Martin Benešík**: Investigation; supervision; methodology; writing – review and editing. **Jitka Krouská**: Investigation; methodology; writing – review and editing. **Petr Sedláček**: Investigation; supervision; writing – review and editing. **Pavel Dvořák**: Conceptualization; funding acquisition; supervision; writing – review and editing.

## Supplementary information

**Fig. S1.**
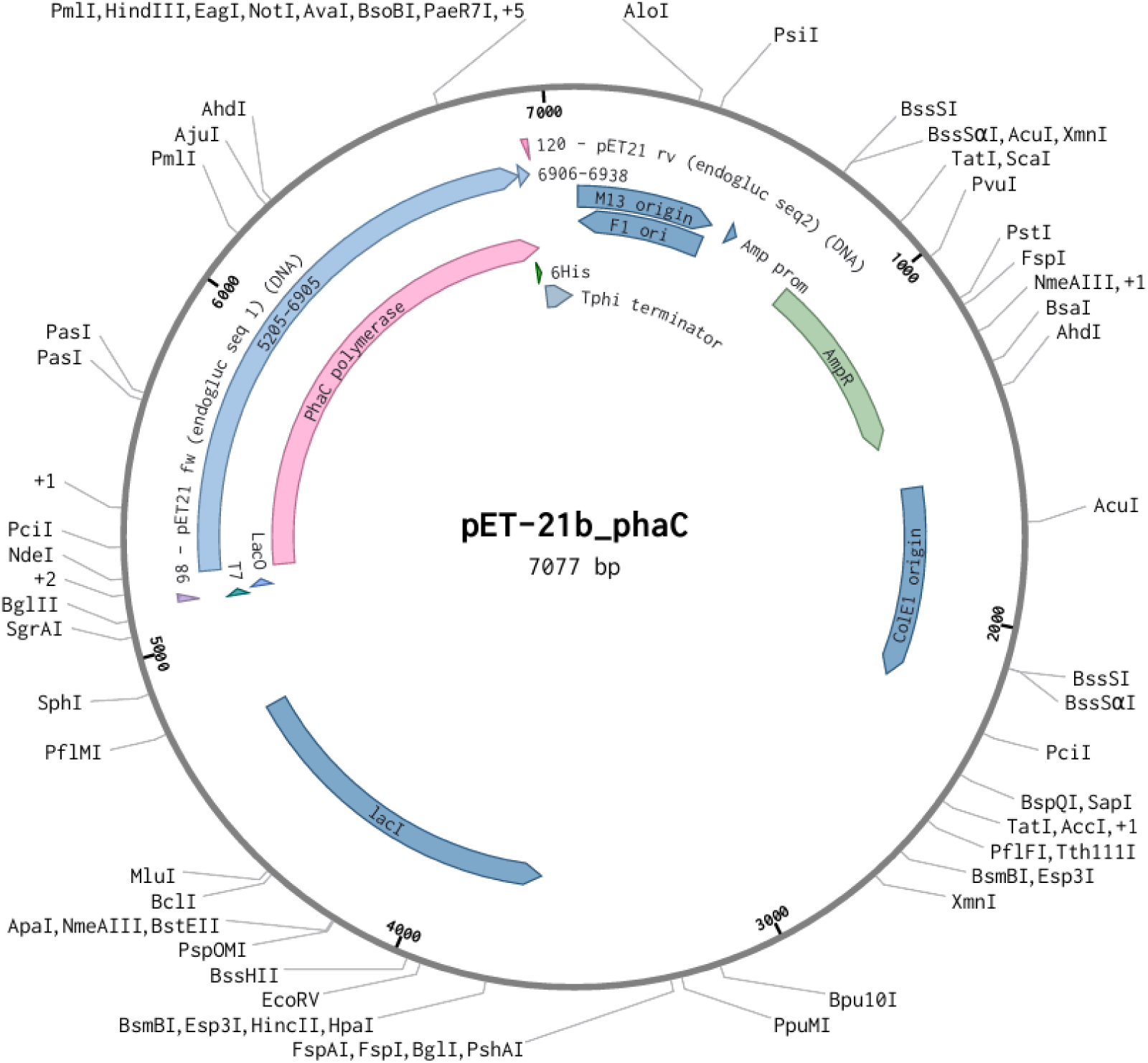
The pET-21b_*phaC* vector map. The scheme of pET-21b vector, containing an ampicillin resistance gene, with the inserted *phaC* gene (locus tag IS481_08630) from *C. thermodepolymerans* DSM15344. The scheme was generated by Benchling.

**Fig. S2.**
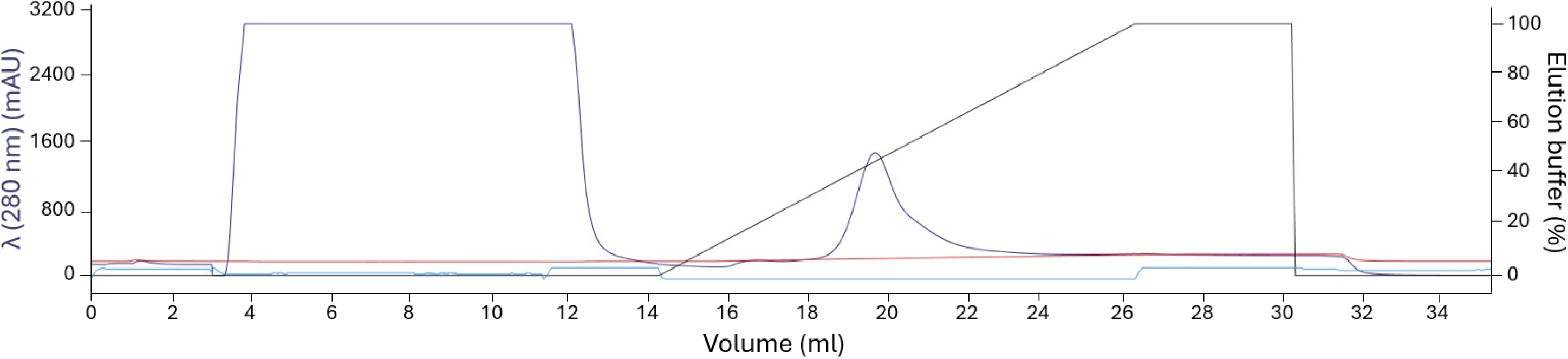
A chromatogram of PhaC purification. Chromatogram from Ni-NTA affinity chromatography (performed using FPLC) of cell-free extract obtained from the *E. coli* BL21(DE3) GOLD cells with pET-21b_*phaC* overproducing PhaC. The protein was eluted in the range of 200–325 mM imidazole. Black line – percentage of elution buffer (imidazole gradient), red line – conductivity, blue line – absorbance at 280 nm, turquoise line – system pressure.

**Fig. S3.**
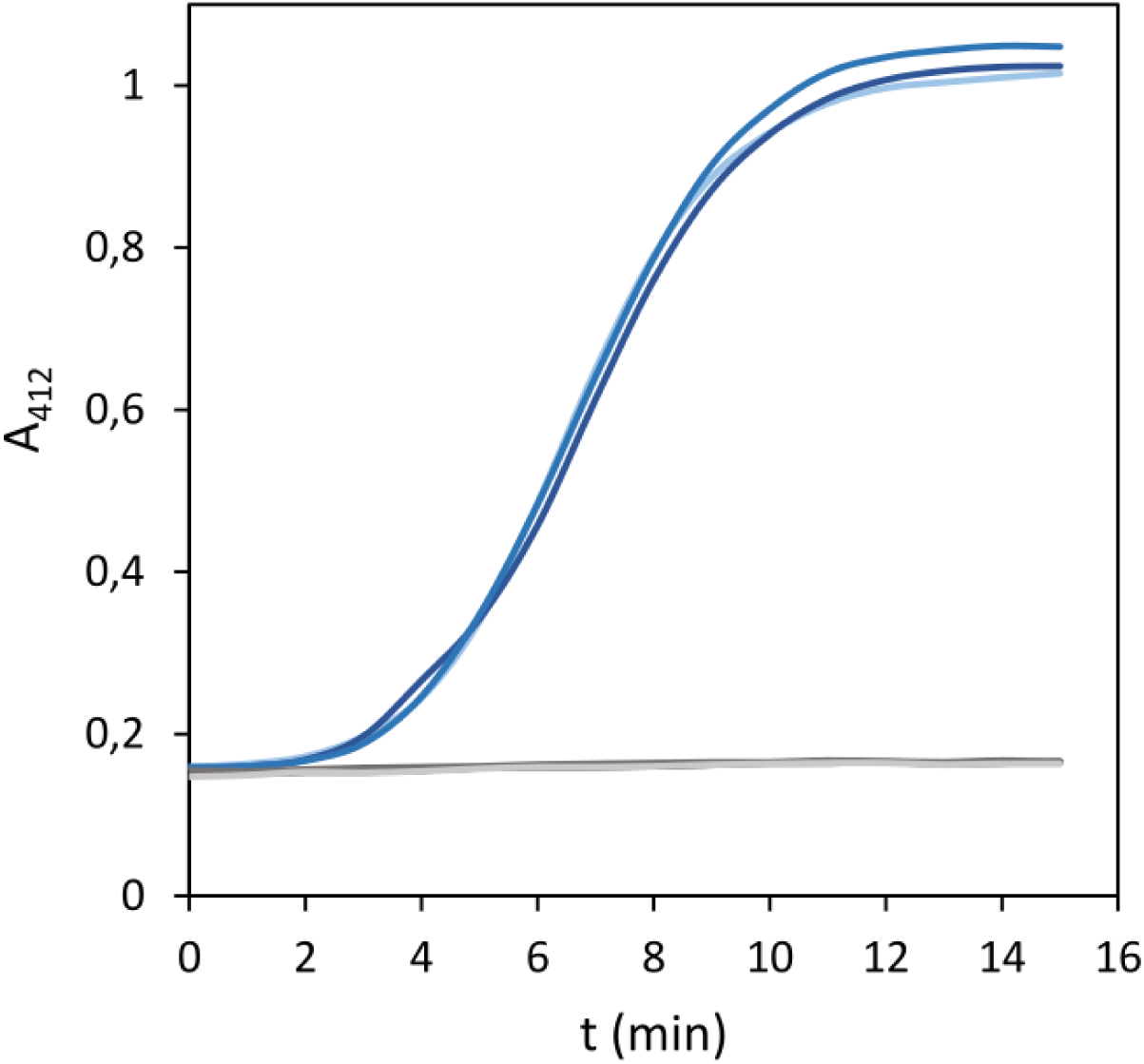
Absorbance change at 412 nm during the PhaC activity measurement. PhaC activity assay, based on Ellman’s reaction, was measured at 50 °C in microplate reader. PhaC (0.5 mg.ml^−1^) polymerise the substrate 3-hydroxybutyryl-CoA and the side product CoA is detected at 412 nm. For negative control PhaC was replaced with Bovine Serum Albumin (0.5 mg.ml^−1^). Blue lines represent PhaC, grey lines represent the negative control. All activity measurements were performed in biological triplicate (n=3) and the mean absorbance changes were used to calculate activity.

**Fig. S4.**
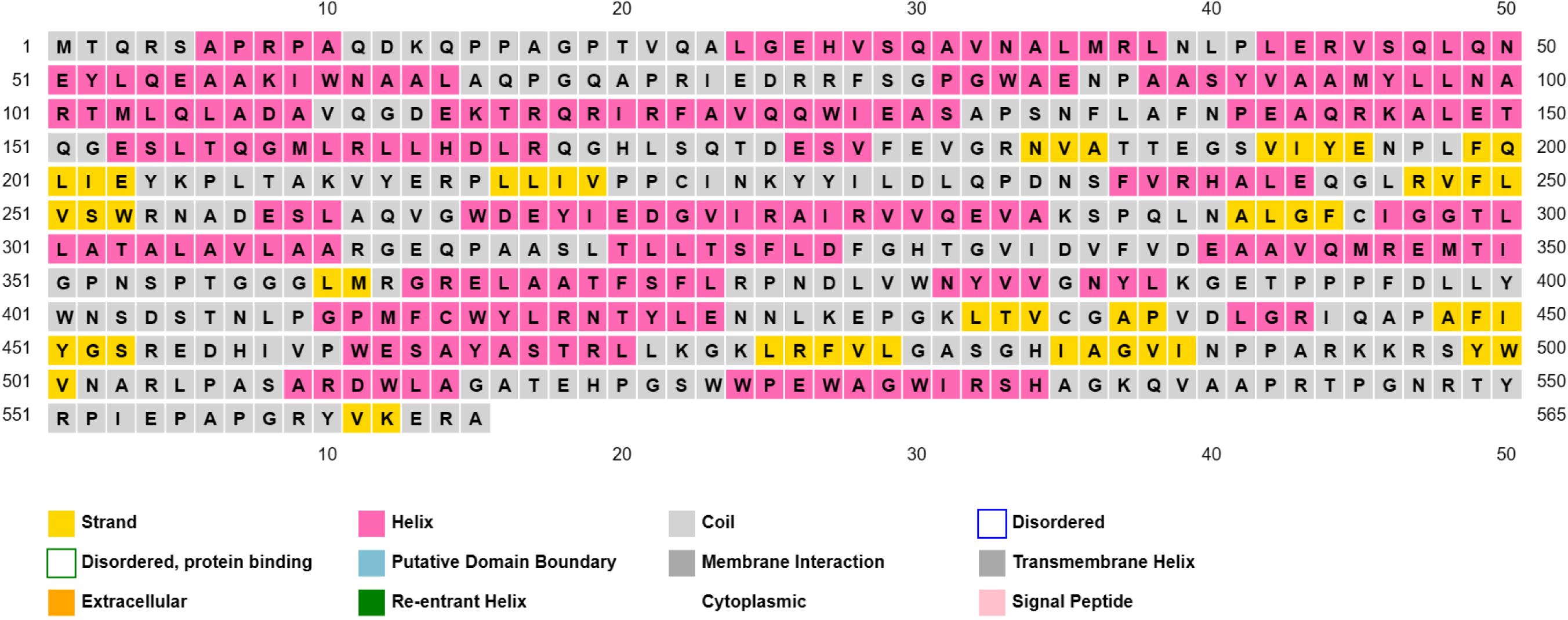
PHA synthase secondary structure scheme generated by the PSIPRED online tool. The α-helices are represented by magenta and β-strands by yellow and coil by grey colour.

**Fig. S5.**
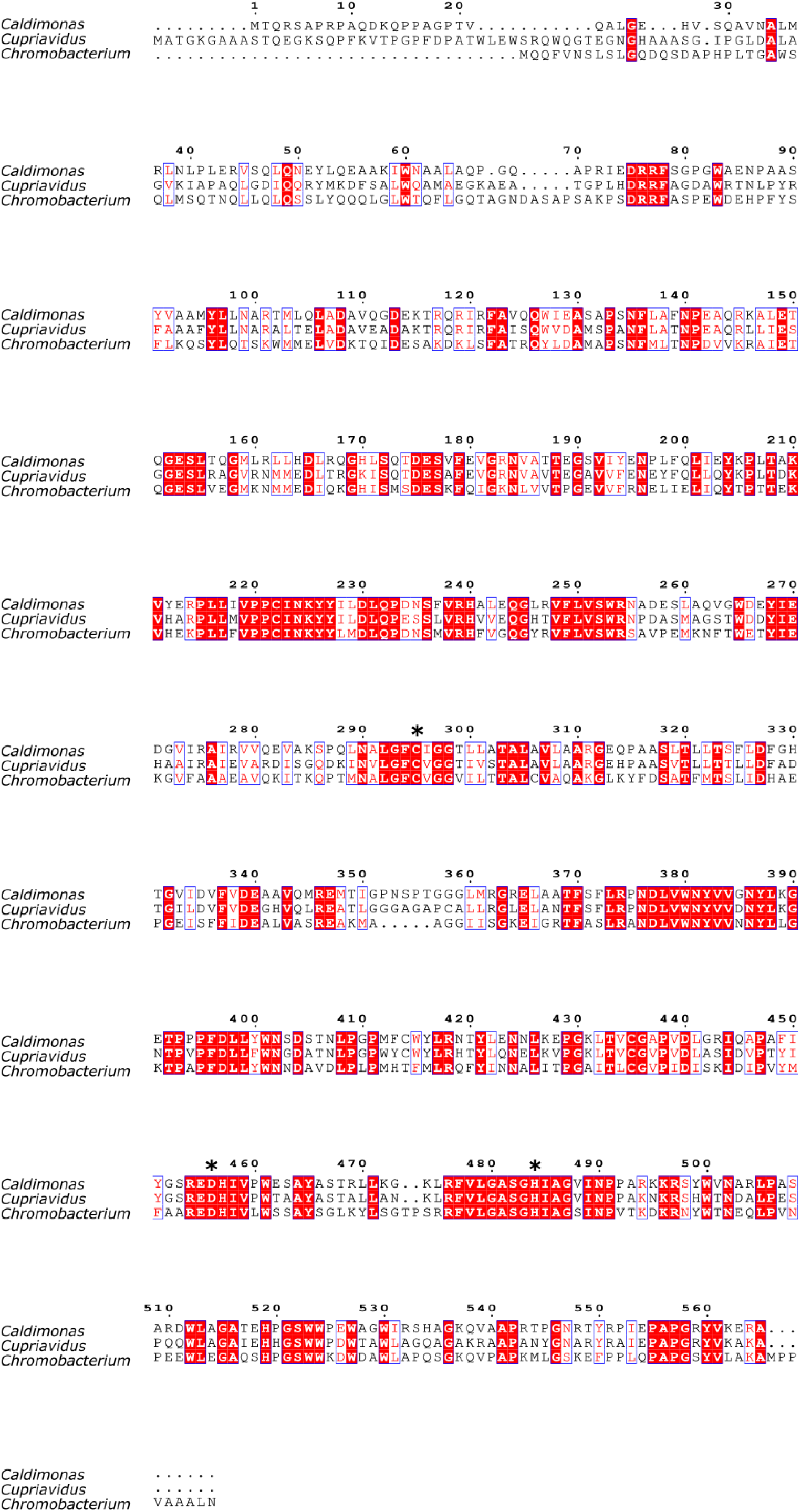
Multiple sequence alignment of PHA synthase protein sequences from *Caldimonas thermodepolymerans, Cupriavidus necator and Chromobacterium sp.* PhaC catalytic triad residues predicted by MSA as C295, D456 and H484 are marked with an asterisk above them. The MSA was performed by Clustal Omega with PhaC sequences from *C. thermodepolymerans* DSM15344, *C. necator* H16 and *Chromobacterium* sp. USM2 and visualised by ESPript 3.2.

**Fig. S6.**
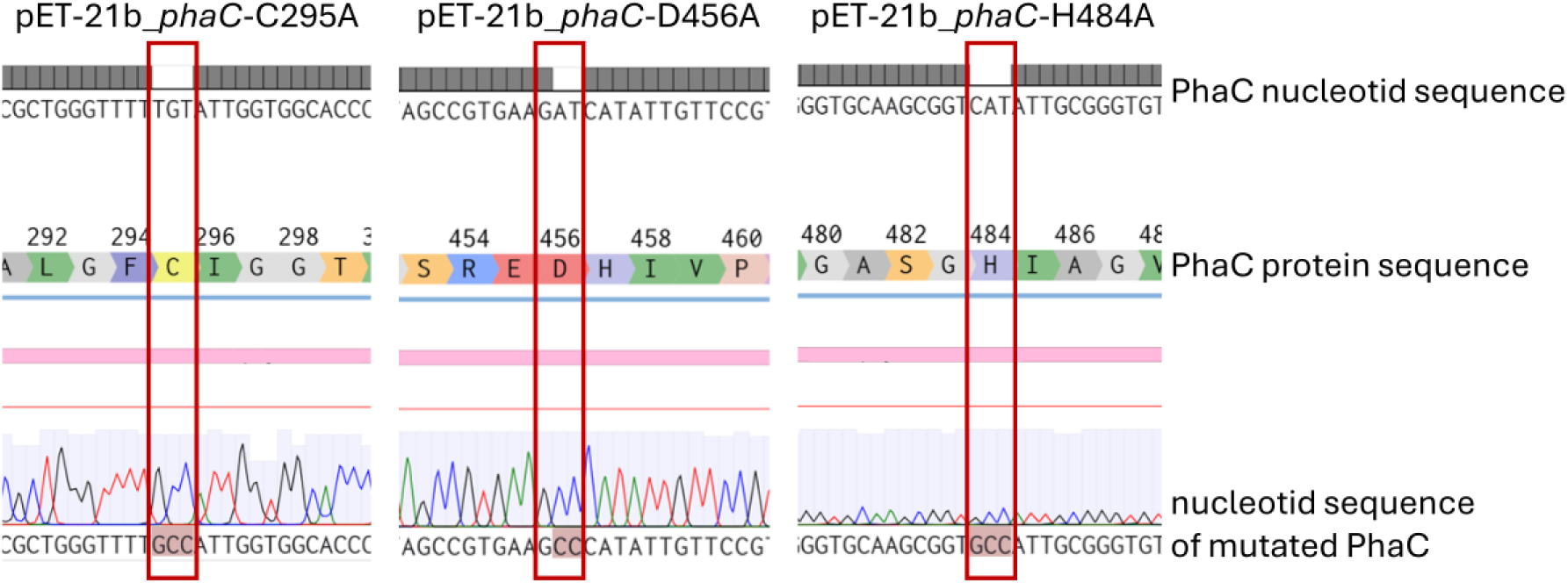
Alignment of *phaC mutant* sequences. All plasmid constructs have successfully mutated codon of the appropriate amino acid to GCC (alanine). Mutated codons are marked with red rectangles.

**Fig. S7.**
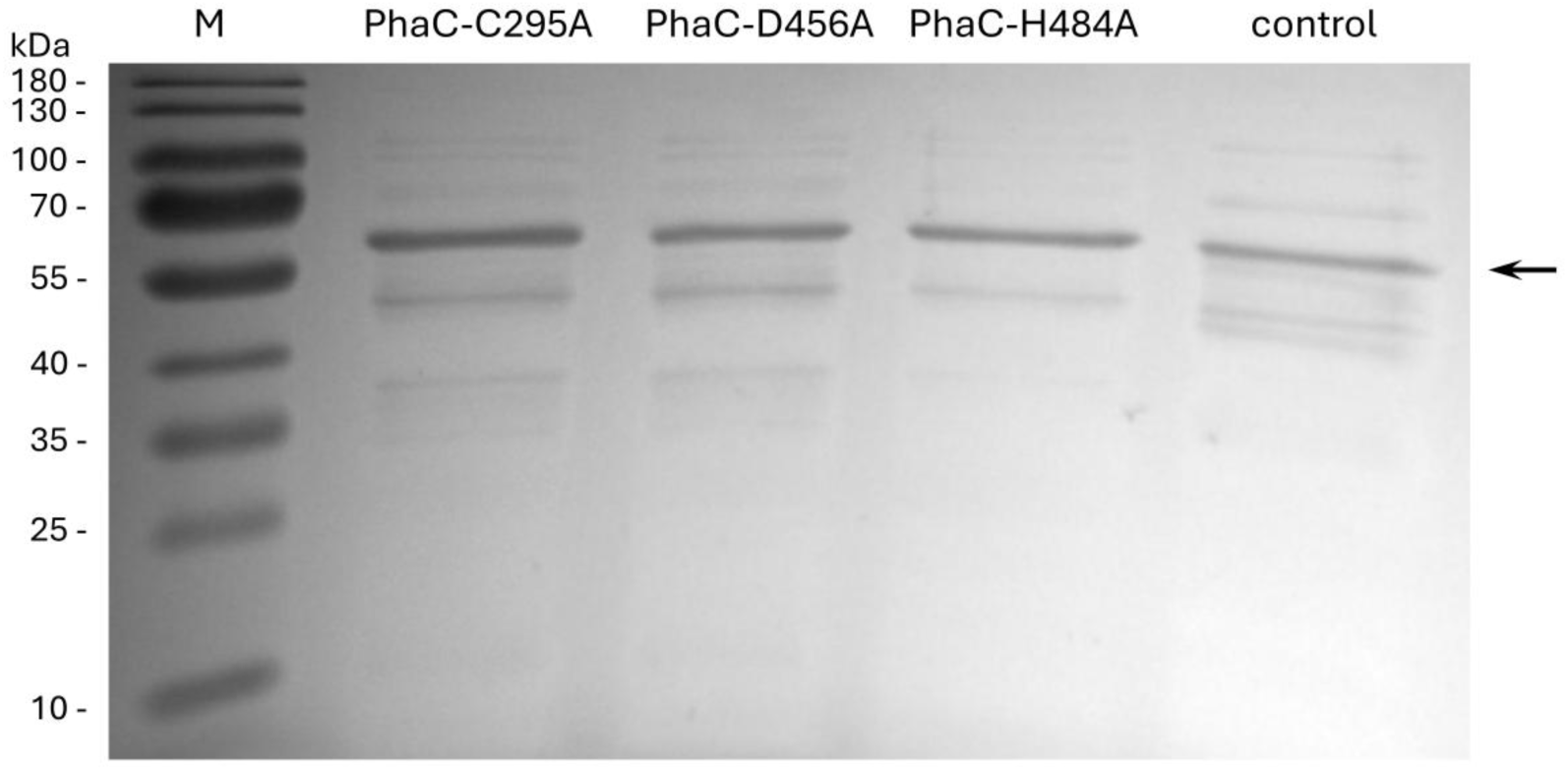
SDS-PAGE of cell-free extracts with overproduced mutated PhaC protein samples before purification. SDS-PAGE analysis showing a band corresponding to the molecular weight of PhaC (62.7 kDa; marked with an arrow) in all three mutants (PhaC-C295A, PhaC-D456A and PhaC-H484A) and also in the negative control (inoculated pET-21b_*phaC*-H484A without IPTG induction). Presence of PhaC in the negative control sample may be caused by an error during cloning, resulting in a non-functional LacI repressor. M – PageRuler Prestained Protein Ladder (10 to 180 kDa).

**Fig. S8.**
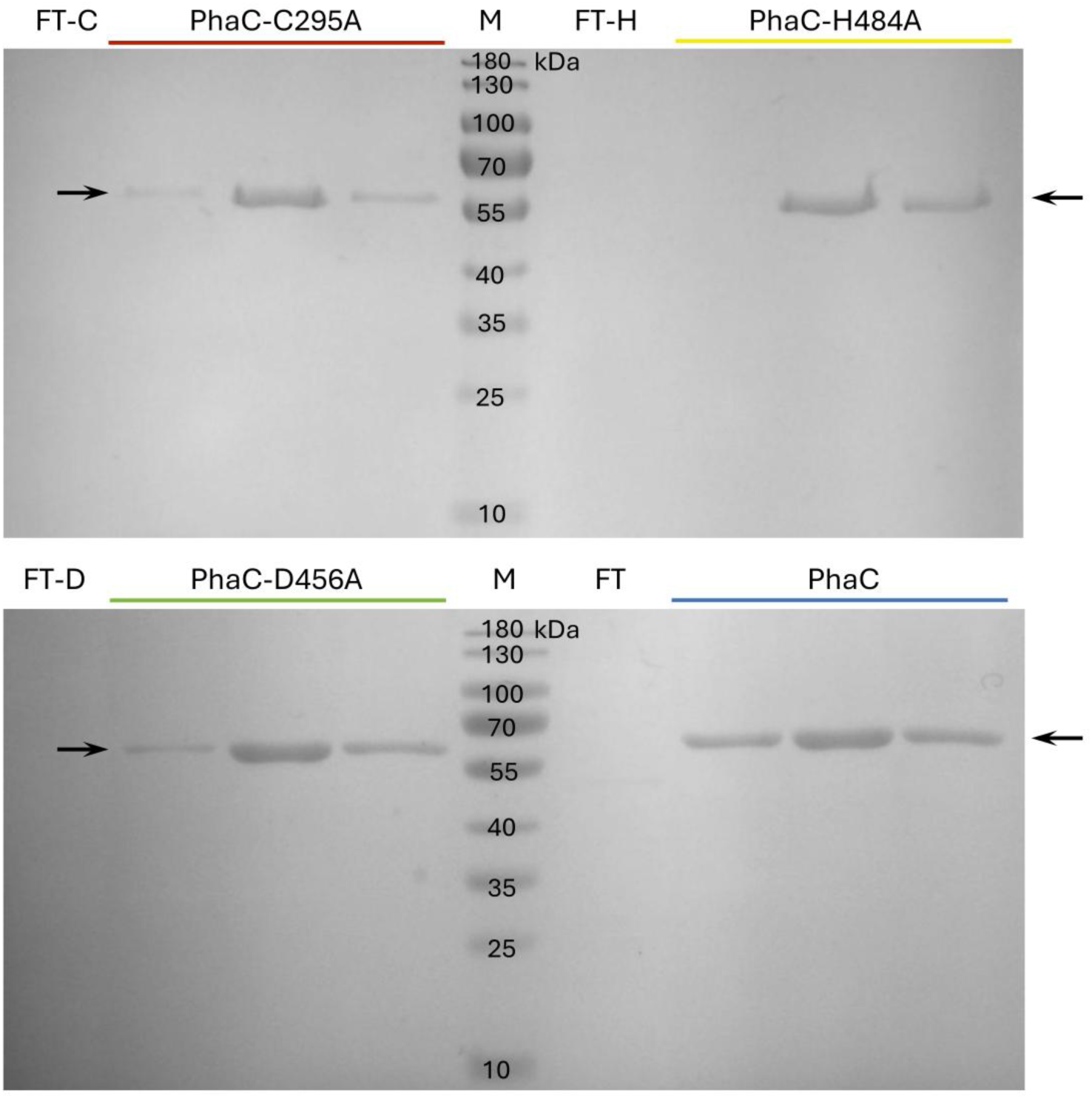
SDS-PAGE of WT PhaC and mutant variants. SDS-PAGE after purification of PhaC and its variants (PhaC-C295A, PhaC-D456A and PhaC-H484A). Purified PhaC and its mutated variants are marked with arrows, FT – flow through, M – PageRuler Prestained Protein Ladder (10 to 180 kDa).

**Table S1.**
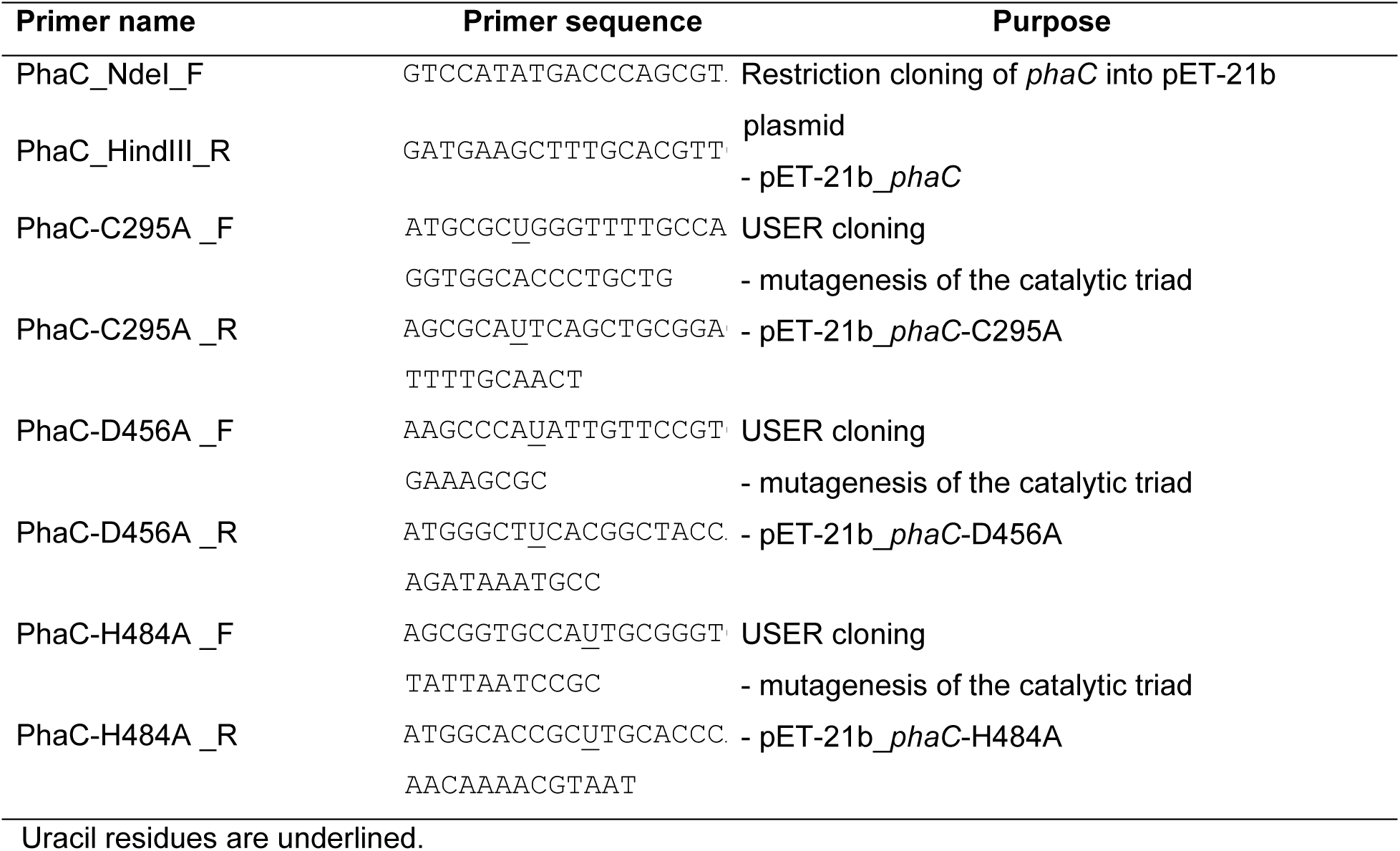
List of primers.

